# Three metabolic pathways replenishing the one-carbon pool collectively support growth and virulence of *Listeria monocytogenes*

**DOI:** 10.1101/2025.09.10.675311

**Authors:** Sandra Freier, Sarah Frentzel, Moritz Müller, Susan Scheffler, Sabrina Wamp, Tim Engelgeh, Janina Döhling, Dunja Bruder, Sascha Kahlfuss, Sven Halbedel

## Abstract

The bacterium *Listeria monocytogenes* can grow in the cytoplasm of infected human cells, but there it relies on specific biosynthetic pathways for intracellular nutrient supply. We previously found that the glycine cleavage system (GCS) is needed for intracellular growth. The GCS decarboxylates glycine for generation of 1C-tetrahydrofolates (1C-THF), folate-dependent one carbon donors needed for biosynthesis of other metabolites. We continued our studies on the GCS and show that a *L. monocytogenes* Δ*gcvPAB* mutant, lacking the GCS glycine dehydrogenase, is attenuated without resembling the phenotype of classical virulence factor mutants. The Δ*gcvPAB* mutant also grew poorly in synthetic medium, explained by the presence of glycine that was toxic for this strain. Selection of glycine resistant suppressors yielded a survivor, in which the N- and C-terminal parts of the formate-tetrahydrofolate ligase (*fhs*) gene, which is naturally separated into two parts by a premature stop codon in the *L. monocytogenes* reference strain EGD-e, were reassembled into a full-length open reading frame. Like the GCS, Fhs also feeds the 1C-THF pool and its restoration cured the virulence defects of the Δ*gcvPAB* mutant. Another suppressor had a mutated *glyA* gene, encoding serine hydroxymethyltransferase, and combinatorial deletions of *gcvPAB* and *glyA* in *fhs^-^* and *fhs*^+^ backgrounds demonstrated a role of GlyA in 1C-THF metabolism. Our results show that three pathways feed the 1C-THF pool to support growth and virulence of *L. monocytogenes* and represent the first example of the spontaneous reactivation of a *L. monocytogenes* gene that is inactivated by a premature stop codon.

## Introduction

*Listeria monocytogenes* is a facultative human pathogen that can cause serious infections after ingestion. To establish an infection, the bacterium first crosses the intestinal-blood barrier by invasion of gut epithelial cells and subsequent transcytosis to the basolateral side of infected cells, where it is released to the bloodstream (Quereda *et al*., 2021). The liver is then the main primary replicative niche of the pathogen, where the bacterium invades hepatocytes, replicates intracellularly and spreads from cell to cell (Koopmans *et al*., 2023). From there, the bacterium can spread hematogenously to the placenta of pregnant women, where it crosses the placental barrier and ultimately causes fetal infections (Charlier *et al*., 2020). *L. monocytogenes* can also invade the brain, possibly achieved by a mechanism similar to that used to cross the intestinal barrier or by infected macrophages that circulate in the blood and are able to cross the blood brain barrier (Disson & Lecuit, 2012). Intra-axonal transport of *L. monocytogenes* from peripheral sites to the brainstem is another proposed mechanism of brain invasion (Bagatella *et al*., 2022).

The bacterium replicates inside the cytoplasm and exploits polymerization of host cell actin to drive intracellular locomotion and spread from cell to cell (Pizarro-Cerda & Cossart, 2018, Quereda *et al*., 2021). To ensure rapid intracellular replication, *L. monocytogenes* has adapted to the specific nutrient availability within the host cell cytoplasm using specific uptake systems such as the hexose phosphate transporter Hpt, enzyme I of the phosphoenolpyruvate:sugar phosphotranferase system, the Opp oligopeptide permease as well as the Cta and Tcy cysteine transporters (Borezee *et al*., 2000, Chico-Calero *et al*., 2002, Xayarath *et al*., 2009, Freeman *et al*., 2025), illustrating the specific importance of certain carbon and nitrogen sources for intracellular nutrition. Likewise, several genes required for the biosynthesis of different cellular building blocks such as purines, aromatic amino acids or menaquinone are required for intracellular replication, while their deletion is tolerated during growth in complex laboratory medium (Stritzker *et al*., 2004, Faith *et al*., 2012, Smith *et al*., 2021, Fischer *et al*., 2022). The conditional essentiality of biosynthetic genes in the host cell cytoplasm shows that certain limitations in the nutrient availability exist intracellularly that *L. monocytogenes* does not encounter in complex laboratory media.

Folate is one of the compounds, which apparently becomes limiting in the host cell as some folate biosynthesis genes are essential for intracellular growth, although they are dispensable during growth in BHI broth (Zhang *et al*., 2022, Feng *et al*., 2023, Stamm *et al*., 2024). The biologically active form of folate is tetrahydrofolate (THF) that transfers one carbon units (1C) to various substrates. 1C-THF species are important for biosynthesis of purines and pyrimidines as well as for the formation of serine, methionine and N-formylmethionine (Green & Matthews, 2007).

Three different pathways ensure 1C-THF formation in *L. monocytogenes*: (i) a two enzyme reaction mediated by the formate-THF ligase Fhs and the bifunctional N5,N10-methylene-THF dehydrogenase/cyclohydrolase FolD (Feng *et al*., 2023), (ii) the serine hydroxymethyltransferase GlyA (Schirch *et al*., 1985), and (iii) the aminomethyltransferase GcvT from the glycine cleavage system (Fujiwara *et al*., 1984) (Fig. 1A). The important role of 1C-THF biosynthesis for intracellular nutrition is reflected by the strong attenuation of a *folD* mutant in macrophages (Feng *et al*., 2023). Likewise, a mutant lacking the *gcvPAB* genes, encoding the two subunits of glycine dehydrogenase, which is another crucial component of the glycine cleavage system, shows impaired replication in macrophages (Fischer *et al*., 2022). The glycine cleavage system (GCS) is a multi-enzyme system that catalyzes the breakdown of glycine, in which the GcvPAB enzyme complex mediates the oxidative decarboxylation of glycine as the first step (Fig. 1A). An important function of this pathway is the generation of N5,N10-methylene-THF to replenish the 1C-THF pool (Kikuchi *et al*., 2008).

**Figure 1:**
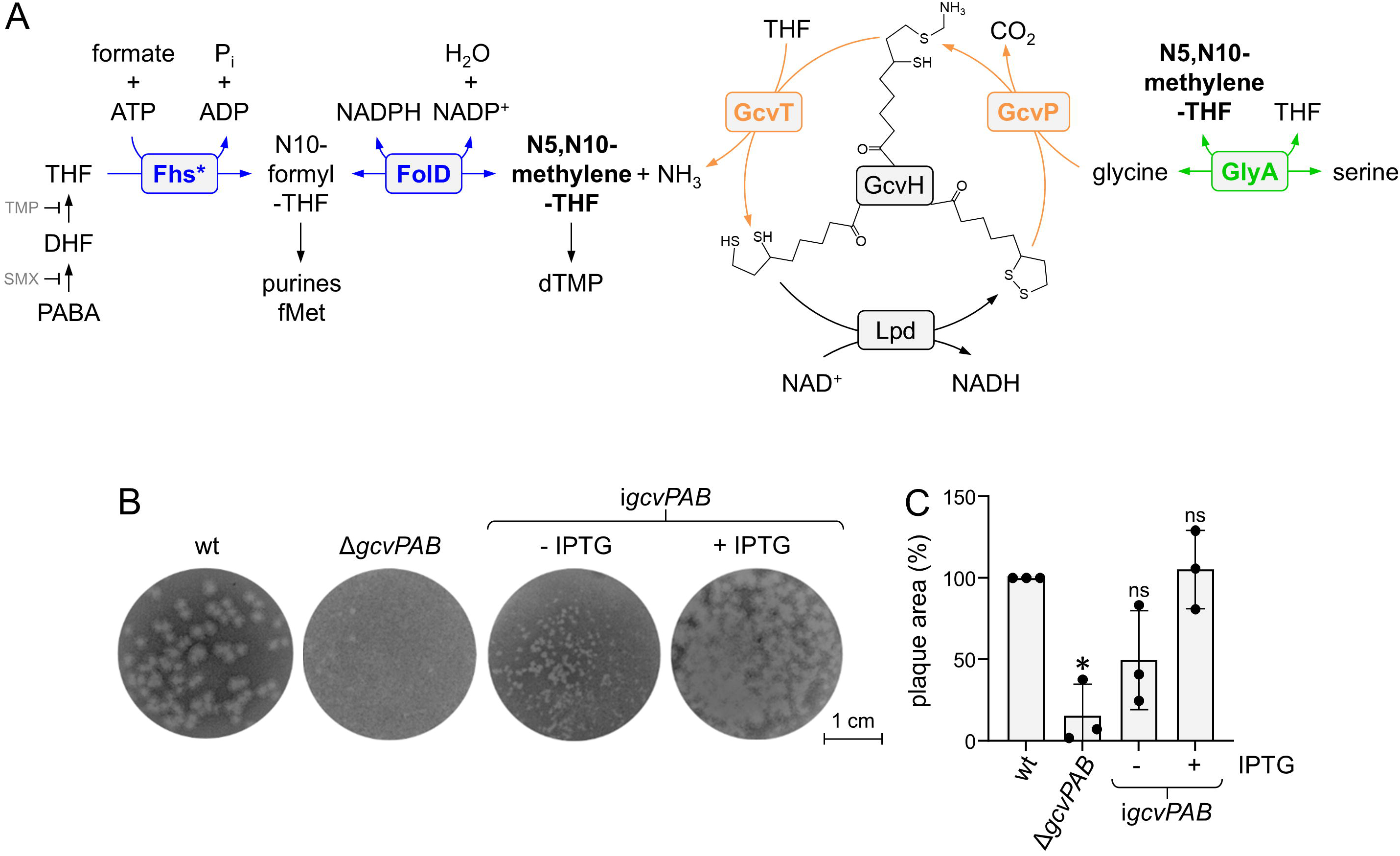
Importance of the Δ*gcvPAB* genes for generation of one carbon donors and for plaque formation in 3T3 mouse fibroblasts. (A) Three enzymatic pathways ensure N5, N10-methylene-THF biosynthesis in *L. monocytogenes*. N5, N10-methylene-THF is generated (i) during conversion of serine to glycine by GlyA (green), (ii) during glycine degradation in the GcvPAB-dependent GCS by GcvP and GcvT (orange) and (iii) by formate THF ligase Fhs in cooperation with the bifunctional N5,N10-methylene-THF dehydrogenase/cyclohydrolase FolD (blue). Biosynthesis of THF from dihydrofolate (DHF) and p-aminobenzoate (PABA) is inhibited by trimethoprim (TMP) and sulfamethoxazole (SMX). (B) Plaque formation assay in 3T3 mouse embryo fibroblasts with *L. monocytogenes* strains EGD-e (wt), LMS305 (Δ*gcvPAB*) and LMS311 (i*gcvPAB*). 1 mM IPTG was added as indicated. (C) Quantification of the assay shown in panel B. Plaque areas were determined using ImageJ and average values and standard deviations were calculated from three independent experiments. Original data points are shown and the asterisk marks a statistically significant difference (*P*<0.01, *t*-test with Bonferroni-Holm correction, ns – not significant).

We here have continued our previously initiated investigation on the attenuated phenotype of a *L. monocytogenes* Δ*gcvPAB* mutant (Fischer *et al*., 2022). Our results indicate that attenuation of this mutant results from limited 1C-THF availability. Isolation of Δ*gcvPAB* suppressor mutants restoring attenuation led to the discovery of a mutation that reactivates the *fhs*/*folD* 1C-THF biosynthesis pathway, which is naturally inactivated by a premature stop codon in the *fhs* gene of *L. monocytogenes* strain EGD-e, a widely used laboratory strain. Furthermore, the three 1C-THF generating pathways were found to be synthetic lethal, further illustrating the importance of folate biosynthesis for growth and virulence of *L. monocytogenes*.

## Results

### In vitro virulence of a L. monocytogenes ΔgcvPAB mutant

We have demonstrated previously that a *L. monocytogenes* Δ*gcvPAB* mutant replicates with reduced growth rate in mouse macrophages and is impaired in cell-to-cell spread in 3T3 mouse fibroblasts (Fischer *et al*., 2022). To further support this observation, we here complemented the Δ*gcvPAB* mutant with an IPTG-inducible *gcvPAB* copy and re-analyzed cell-to-cell spread. In agreement with our previous results, only small plaques were formed in 3T3 cells upon infection with the Δ*gcvPAB* mutant (plaque area: 15±19% of wild type level) and small plaques were also formed by the complemented strain in the absence of IPTG (50±19%). However, plaque formation was restored when IPTG was added (105±24%, Fig. 1B-C), as expected.

To further study this virulence defect, dissemination of a Δ*gcvPAB* strain expressing the red fluorescent protein DsRed-Express in infected 3T3 cultures was analyzed microscopically. The wild type and the Δ*gcvPAB* mutant were found disseminated throughout the cytoplasm of infected cells and neighbor cells also contained bacteria (Fig. S1). In contrast, the Δ*actA* mutant, which cannot spread due to the absence of the actin tail nucleating ActA protein (Kocks *et al*., 1992), formed concentrated foci of fluorescent bacteria in the cytoplasm of infected cells and neighbor cells were usually not infected (Fig. S1). As the phenotypes of the Δ*gcvPAB* and Δ*actA* mutants were different in this assay, the plaque formation defect of the Δ*gcvPAB* mutant must either be caused by an inability to invade the cells or to replicate within them. To discriminate between these possibilities, we next quantified invasion and intracellular replication of the Δ*gcvPAB* mutant in comparison to well characterized mutants lacking either the *hly* or *actA* genes in 3T3 cells. As can be seen in Fig. 2A, invasion into 3T3 cells was not impaired, however, intracellular growth was retarded compared to wild type and to the Δ*actA* mutant, which cannot spread at all but otherwise grows normally (Kocks *et al*., 1992). Similarly, intracellular growth of the Δ*gcvPAB* mutant was delayed in J774 macrophages (Fig. 2B), however, no delay was detected in HepG2 hepatocytes (Fig. 2C). The Δ*gcvPAB* mutant was as hemolytic (Fig. 2D-E) and as resistant against lysozyme as the wild type (Fig. 2F). Differences in hydrogen peroxide sensitivity were also not found as the minimal inhibitory H2O2 concentrations was 3.1 mM for both strains. Taken together, the Δ*gcvPAB* mutant shows delayed intracellular growth in fibroblasts, explaining the plaque formation defect, and the delayed growth of the Δ*gcvPAB* mutant observed in macrophages is not related to common pathogen defense strategies of macrophages.

**Figure 2:**
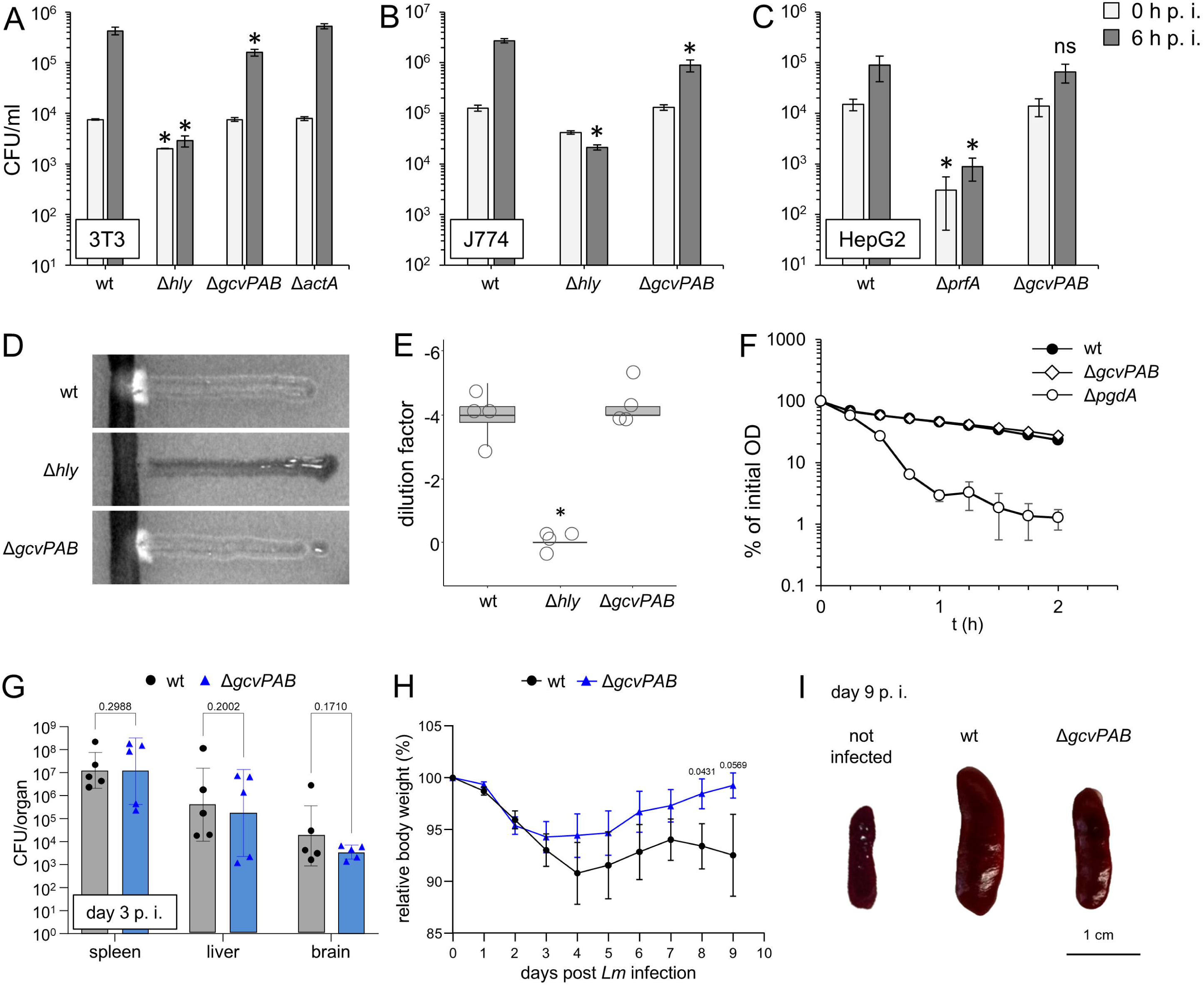
*In vitro* and *in vivo* virulence of the Δ*gcvPAB* mutant. (A) Replication of *L. monocytogenes* strains EGD-e (wt), LMS305 (Δ*gcvPAB*), LMS250 (Δ*hly*, replication deficient control) amd LMS251 (Δ*actA*, spreading deficient control) in 3T3 mouse fibroblasts. (B) Replication of *L. monocytogenes* strains EGD-e (wt), LMS305 (Δ*gcvPAB*) and LMS250 (Δ*hly*, negative control) in J774 mouse macrophages. (C) Replication of *L. monocytogenes* strains EGD-e (wt), LMS305 (Δ*gcvPAB*) and BUG2214 (Δ*prfA*, invasion and replication deficient control) in HepG2 human hepatocytes. The experiments shown in Figures A–C were repeated three times, with each run consisting of technical replicates (n = 3). The mean values and standard deviations were calculated from the technical replicates of a representative run. Asterisks mark statistically significant differences (panels A and B: *P*<0.01 *t*-test with Bonferroni-Holm correction, panel C: *P*<0.05 *t*-test). (D) CAMP assay to compare hemolysis in *L. monocytogenes* strains EGD-e (wt), LMS305 (Δ*gcvPAB*) and LMS250 (Δ*hly*, negative control). (E) Quantification of hemolysis activity in the same set of strains towards human erythrocytes. Hemolysis activity is expressed as the number of ten-fold dilutions of the various culture supernatants after which no hemolysis could be observed anymore. The experiment was carried out four times. The asterisk marks a statistically significant difference (*P*<0.01, *t*-test with Bonferroni-Holm correction). (F) Lysozyme-induced lysis of *L. monocytogenes* strains EGD-e (wt), LMS305 (Δ*gcvPAB*) and LMS163 (Δ*pgdA*, positive control). The experiment was repeated three times, with each run consisting of technical replicates (n = 3). Mean values and standard deviations were calculated from the technical replicates of a representative run. (G-I) Virulence of the Δ*gcvPAB* mutant in mice. Infection was conducted with 1-20 x 10^4^ CFUs/ml injected into the tail vein of the mice. Five mice were infected with either EGD-e or the Δ*gcvPAB* mutant. (G) Three days post-infection, CFU were quantified in the spleen, liver and brain to determine the bacterial burden. The geometric mean with the geometric standard deviation (SD) is illustrated. (H) Following infection, the mice were scored on a daily basis for weight loss as a parameter of disease severity during *L. monocytogenes* infection. The body weight is presented in relation to the weight prior to infection. The standard error of the mean (SEM) is shown. One representative experiment out of two independent repetitions is shown. (I) Nine days post-infection, the spleens were isolated and compared for organ size under infections with different *L. monocytogenes* strains. Statistical analysis was conducted utilising the GraphPad Prism software, performing unpaired *t*-tests (G – two-tailed, H – one-tailed).

### In vivo virulence of the L. monocytogenes ΔgcvPAB mutant

In order to confirm the virulence attenuation of the Δ*gcvPAB* mutant *in vivo*, C57BL/6J mice were infected intravenously (i.v.) with the EGD-e or Δ*gcvPAB* strains. At day 3 post infection (p.i.), we determined the CFUs within different organs (Fig. 2G). Here, we detected a trend of reduced CFU numbers in the spleen, liver and brain from mice that were infected with the Δ*gcvPAB* mutant, but this did not reach statistical significance. Importantly, however, mice infected with the EGD-e strain showed a more severe body weight loss compared to mice infected with the Δ*gcvPAB* mutant (Fig. 2H). In line with this, the more pronounced enlargement of the spleens observed in mice infected with EGD-e compared to those infected with the Δ*gcvPAB* mutant at day 9 p. i. indicated a reduced pathogenicity of the Δ*gcvPAB* mutant strain (Fig. 2I). Together, our data indicate a moderately ameliorated disease progression in mice infected with the Δ*gcvPAB* mutant compared to mice infected with the EGD-e reference strain.

### Growth defect of a *L. monocytogenes* Δ*gcvPAB* mutant in synthetic medium

To further investigate the phenotype of the Δ*gcvPAB* mutant, we aimed at the identification of extracellular growth conditions that would mimic the intracellular growth defect. In the course of this search we noticed that the Δ*gcvPAB* mutant had a remarkable growth defect in *Listeria* synthetic medium (LSM). LSM broth is a chemically defined medium that contains all components required for growth at defined concentrations, but it has not been chemically validated to reflect host cytosolic conditions (Whiteley *et al*., 2017). When cultivated in LSM broth at 37°C, growth of the Δ*gcvPAB* mutant was strongly retarded, which was in stark in contrast to BHI broth, where no growth defect was apparent (Fig. 3A). This growth defect was complemented as strain LMS311 carrying an IPTG-inducible *gcvPAB* copy grew as slow as the parental mutant in the absence and as fast as the wild type in the presence of IPTG (Fig. 3B). This demonstrates that the decarboxylase component of the glycine cleavage system is also required for normal growth in synthetic LSM medium.

**Figure 3:**
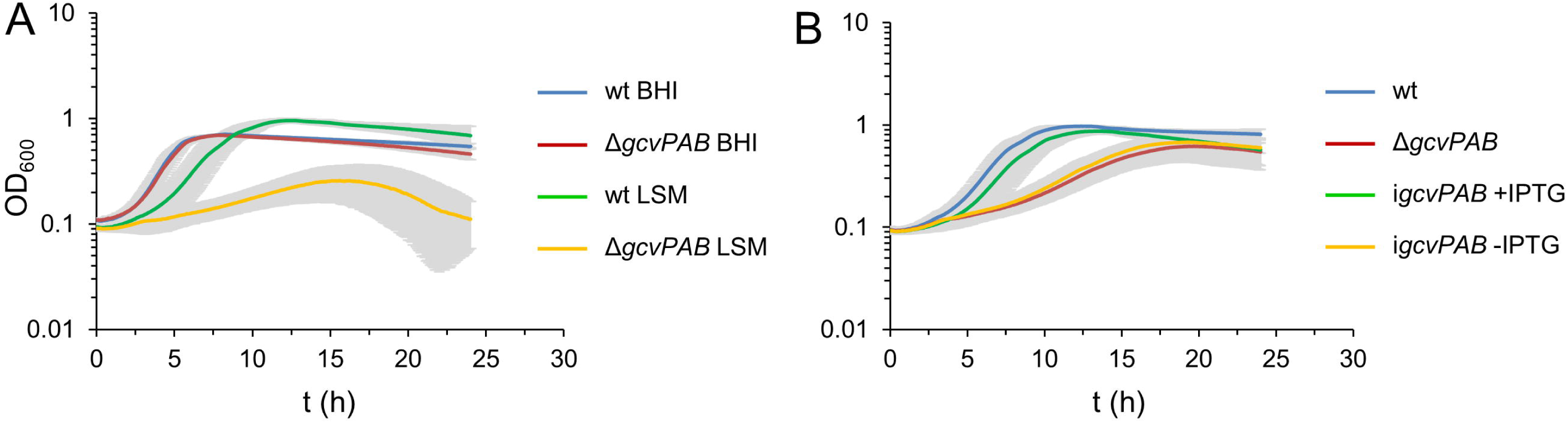
Growth of the Δ*gcvPAB* mutant in laboratory media. (A) Growth of *L. monocytogenes* strains EGD-e (wt) and LMS305 (Δ*gcvPAB*) in complex BHI and chemically defined LSM medium. (B) Complementation of the growth defect of Δ*gcvPAB* mutant in LSM medium. Growth of *L. monocytogenes* strains EGD-e (wt), LMS305 (Δ*gcvPAB*) and LMS311 (i*gcvPAB*) in LSM medium ± 1 mM IPTG. The experiment was repeated three times, with each run consisting of technical replicates (n = 3). Mean values and standard deviations were calculated from the technical replicates of a representative run.

### High glycine concentrations are toxic for the Δ*gcvPAB* mutant

Breakdown of glycine in the GCS generates 1C-THF, which serves as an important one carbon unit donor in various biosynthesis pathways (Fig. 1A). If glycine cannot be catabolized (and 1C-THF cannot be generated) by the GCS due to deletion of *gcvPAB*, glycine might be re-routed to the serine hydroxymethyl transferase GlyA for serine formation, even though this would consume 1C-THF and therefore even further deplete the cell for 1C-THF. We therefore considered the possibility that glycine might become toxic in the absence of the GCS as observed in a GCS mutant of the cyanobacterium *Synechocystis* (Eisenhut *et al*., 2007) due to depletion of 1C-THF pool. To test this, we determined the growth of the wild type and the Δ*gcvPAB* mutant in the presence of varying glycine concentrations. As can be seen in Fig. 4A, the wild type was able to grow without glycine and a tenfold increase of the glycine concentration also had no effect. Apparently, glycine can be generated, presumably from serine (which is present in LSM broth) via GlyA, when it is not supplied externally, and does not become toxic in the presence of a functional GCS. In contrast, the Δ*gcvPAB* mutant exhibited delayed growth in LSM with standard glycine concentration, but growth was accelerated when the glycine concentration was halved and even reached wild type level, when it was further reduced (Fig. 4B). In the complete absence of glycine, growth of the Δ*gcvPAB* mutant was largely unaffected, presumably because glycine cannot be converted to serine by GlyA anymore, thereby conserving the 1C-THF pool. In contrast, a tenfold increase in glycine concentration significantly impaired growth. (Fig. 4B), whereas alterations in the serine concentrations had no effect on the growth of either strain (Fig. S2). This demonstrates that glycine or a metabolite of glycine becomes toxic in the absence of a functional GCS. That the Δ*gcvPAB* mutant can even grow without glycine also reinforces the idea that glycine must be made from serine through GlyA, since glycine formation through the GCS running in a reverse reductive mode would not be possible in the absence of the GcvP component (Yishai *et al*., 2018).

**Figure 4:**
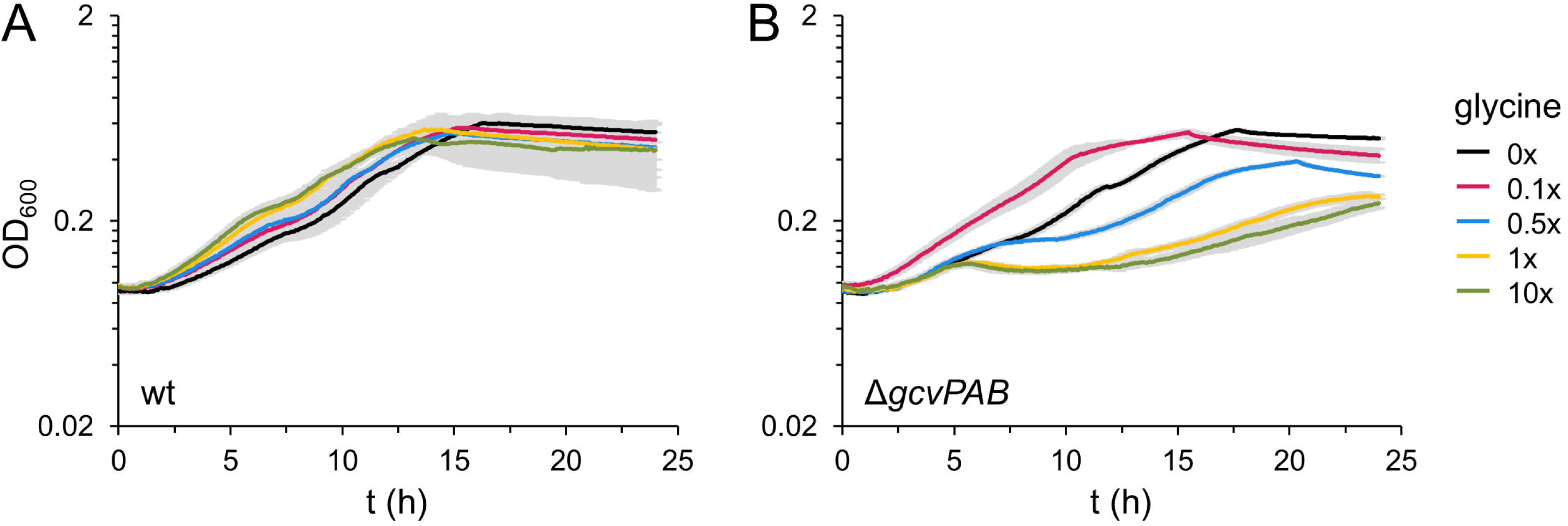
The Δ*gcvPAB* mutant is sensitive to increased glycine concentrations. (A-B) Growth of *L. monocytogenes* strains EGD-e (wt, A) and LMS305 (Δ*gcvPAB*, B) in LSM medium containing different glycine concentrations. Glycine concentrations are expressed relative to standard LSM concentrations (1x = 1.3 mM). The experiment was repeated three times, with each run consisting of technical replicates (n = 3). Mean values and standard deviations were calculated from the technical replicates of a representative run.

### Mutations suppressing glycine sensitivity of the Δ*gcvPAB* mutant

We next exploited glycine sensitivity to screen for mutants suppressing the Δ*gcvPAB* phenotype. For this, the Δ*gcvPAB* mutant was cultivated in LSM medium containing 100-fold the amount of glycine as present in the standard recipe. Cells were grown for 24 hours and then plated on LSM plates containing 100-fold the standard amount of glycine, a condition under which the Δ*gcvPAB* mutant would not grow. Colonies that could grow on these plates were isolated and their genomes were sequenced. Glycine insensitive suppressors carried mutations in the *codY* (*lmo1280*), *fhs* (*lmo1877*), *folK* (*lmo0226*) and *glyA* (*lmo2539*) genes. Suppressor LMSF3 carries a G236E substitution in the *codY* gene encoding the transcriptional repressor of the CodY regulon. LMSF8 had a G82R exchange in *folK* coding for 7,8-dihydro-6-hydroxymethylpterin pyrophosphokinase that acts in folate biosynthesis. LMSF10 had a G62S substitution in *glyA* (encoding the gene for serine hydroxymethyltransferase for serine/glycine interconversion) in addition to the *codY* G236E exchange that is also present in LMSF3. However, the most remarkable suppressor mutation was found in strain LMSF15, in which the N-and C-terminal parts of the *fhs* pseudogenes that are separated by a premature stop codon and thus inactivated in EGD-e are reunited by the introduction of four nucleotides at the end of the N-terminal *fhs* pseudogene *lmo1877*. Here, introduction of a quadruplet (GTGG) restores the complete *fhs* reading frame as it is found in strain 10403S but with one extra valine inserted at the fusion site (Fig. 5A). All four suppressors grew as the wild type in BHI and LSM broth without glycine. However, all suppressors grew better than the parental Δ*gcvPAB* strain in LSM broth containing 100-fold the glycine concentration of the standard recipe. None of the suppressors restored wild type-like growth, but restoration of full length *fhs* (named *fhs^+^* here) had the strongest effect (Fig. 5B).

**Figure 5:**
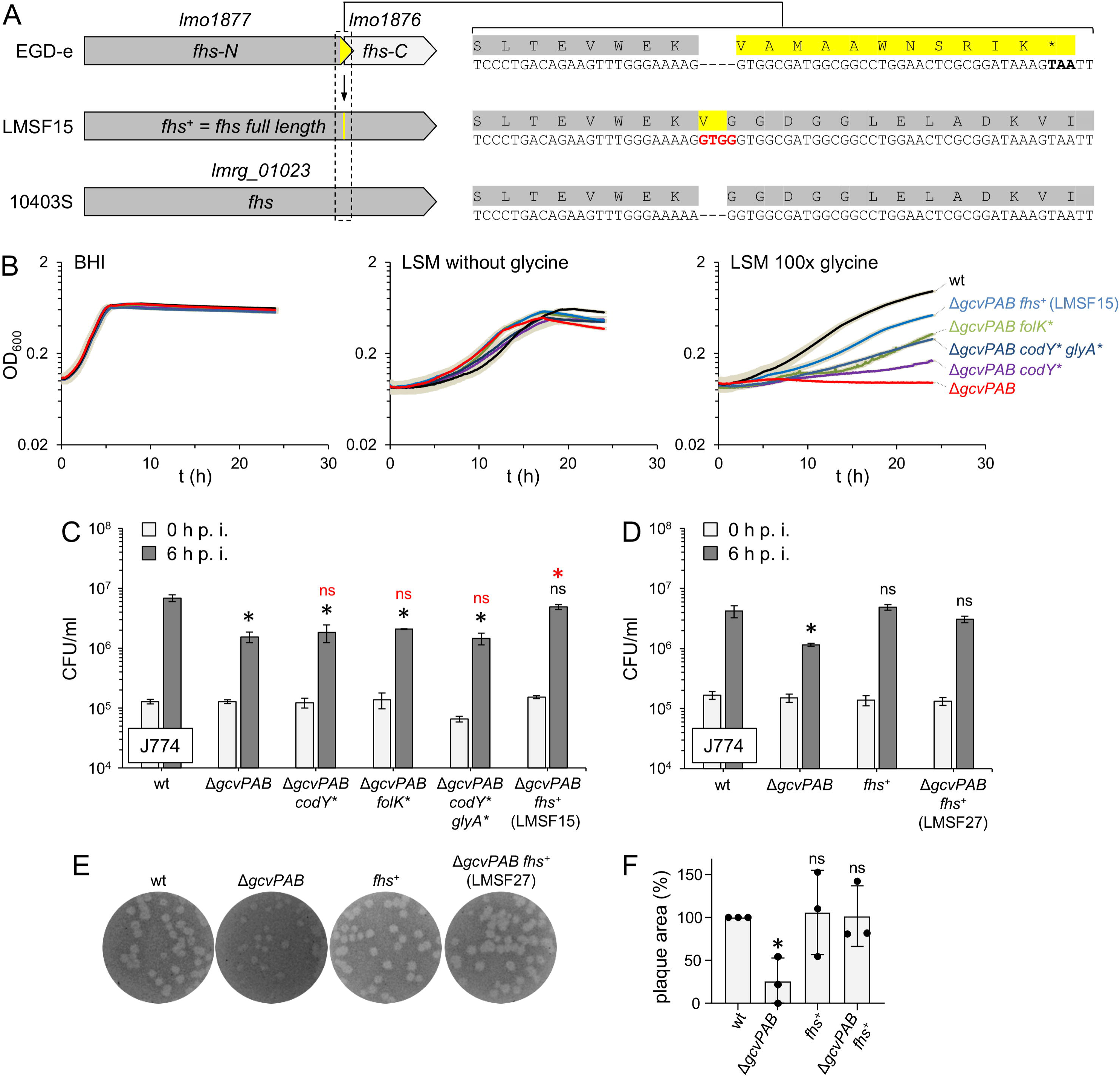
Suppression of the Δ*gcvPAB* phenotype by a *fhs^+^* mutation restoring formate-tetrahydrofolate ligase activity. (A) Restoration of the full length *fhs* open reading frame in Δ*gcvPAB* suppressor strain LMSF15. Schematic illustration of the *fhs* loci in the two reference strains 10403S (full length) and EGD-e (split into two pseudogenes due to a frameshift mutation in *lmo1877*) as well as in the Δ*gcvPAB* suppressor strain LMSF15 where an GTGG insertion (red) restores the full length *fhs* gene. (B) Growth of Δ*gcvPAB* suppressor strains in various media. Strains EGD-e (wt), LMS305 (Δ*gcvPAB*), LMSF3 (Δ*gcvPAB codY G236E*), LMSF8 (Δ*gcvPAB folK G82R*), LMSF10 (Δ*gcvPAB codY G236E glyA G62S*) and LMSF15 (Δ*gcvPAB fhs^+^*) were grown in BHI broth (left panel), LSM medium without glycine (middle panel) and LSM medium supplemented with 100-fold the amount of glycine than in the standard recipe (right panel). Growth curves were repeated three times, with each run consisting of technical replicates (n = 3). Mean values and standard deviations were calculated from the technical replicates of a representative run. (C) Intracellular growth of Δ*gcvPAB* suppressors in J774 mouse macrophages. The same set of strains as in the panel B was used to infect J774 macrophages and the bacterial load six hours post infection (p. i.) was determined. Infections were repeated three times, with each repetition consisting of technical replicates (n = 3). Mean values and standard deviations were calculated from the technical replicates of a representative experiment. Asterisks mark statistically significant differences compared to wild type (black) or compared to the Δ*gcvPAB* mutant (red, *P*<0.01, *t*-test with Bonferroni-Holm correction, ns – not significant). (D) Recreation of the *fhs*^+^ mutation in the Δ*gcvPAB* background confirms suppression of Δ*gcvPAB in vitro* virulence phenotypes by restoration of Fhs activity. Intracellular replication of *L. monocytogenes* strains EGD-e (wt), LMS305 (Δ*gcvPAB*), LMSF26 (*fhs^+^*) and LMSF27 (Δ*gcvPAB fhs^+^*) in J774 mouse macrophages. Strains LMSF26 and LMSF27 were generated from EGD-e and LMS305, respectively, by introduction of the isolated *fhs*^+^ mutation. Infection experiments were repeated three times, with each repetition consisting of technical replicates (n = 3). Mean values and standard deviations were calculated from the technical replicates of a representative experiment. Asterisks mark statistically significant differences compared to wild type (*P*<0.05, *t*-test with Bonferroni-Holm correction, ns = not significant). (E) Plaque formation in 3T3 mouse fibroblasts of the same set of strains as in panel D. (F) Quantification of the assay shown in panel E. Plaque areas were determined using ImageJ and average values and standard deviations were calculated from three independent experiments. Original data points are shown and the asterisk marks a statistically significant difference (*P*<0.01, *t*-test with Bonferroni-Holm correction, ns – not significant).

### Specific suppression of the Δ*gcvPAB* virulence defects by Fhs restoration

In order to determine to what degree suppression of glycine toxicity also repairs the intracellular growth defect in the four suppressor mutants, their growth inside J774 cells was measured. Growth of the Δ*gcvPAB* suppressors with mutations in *codY*, *folK* and *glyA* was as retarded as observed in the parental Δ*gcvPAB* mutant. However, restoration of full length *fhs* suppressed this defect and the Δ*gcvPAB* suppressor with the restored *fhs* gene grew as fast the wild type (Fig. 5C). To further confirm these observations, we generated a plasmid that allows restoration of full length *fhs* in strain EGD-e and its descendants. Using this plasmid, full length *fhs* was generated in the Δ*gcvPAB* mutant and the growth of the resulting strain in mouse macrophages was determined. As can be seen in Fig. 5D, *fhs* repair in the Δ*gcvPAB* background restored wild-type like intracellular replication. Moreover, the Δ*gcvPAB* mutant with the repaired *fhs* gene also formed plaques as the wild type (Fig. 5E-F). The *fhs* gene was also repaired in EGD-e, but this did not further enhance plaquing efficiency or intracellular replication (Fig. 5D-F).

### Frequency of *fhs* truncations in other *L. monocytogenes* strains

Given the important role of *fhs* for growth and virulence of the EGD-e reference strain, we investigated how frequently premature stop codons leading to *fhs* inactivation occur among *L. monocytogenes* isolates from clinical, food, and environmental sources. To this end, we analyzed 29,094 phylogenetically diverse *L. monocytogenes* genomes deposited in the NCBI Pathogen Detection database between 2010 and 2019. The *fhs* loci were extracted by allele calling using MBioSEQ Ridom SeqSphere+ (Bruker), with the *fhs* open reading frame of strain 10403S (*lmrg_01023*) serving as the reference sequence. The extracted sequences were subsequently screened for premature stop codons and frameshift mutations.

Our analysis revealed that the vast majority of strains carried full-length *fhs* genes. Truncated *fhs* variants were detected only in EGD-e and four of its descendants, as well as in five sequence type (ST) ST2 strains belonging to the same SNP cluster (PDS000024994.1), which had been isolated from cheese between 2016 and 2017. These findings indicate that *fhs* inactivation due to premature stop codons is a rare event in *L. monocytogenes*, but that it has arisen independently at least twice during the evolutionary history of the species.

### Three 1C-THF generating pathways support growth, virulence and purine biosynthesis

To compare the contribution of the three 1C-THF generating pathways, *i. e.* the GCS, GlyA and the Fhs/FolD pathway, to growth of *L. monocytogenes*, we sought to generate mutants lacking all three pathways simultaneously. For this, a Δ*glyA* mutant was constructed in EGD-e first. The Δ*glyA* mutant could not grow in LSM broth not containing glycine. However, normal growth was observed, when LSM was supplemented with high glycine concentrations (Fig. S3A-B). Thus, GlyA is needed for glycine generation from serine when glycine is not supplied externally. In contrast, the Δ*glyA* mutant could grow normally in the absence of serine (Fig. S3C-D), which showed that serine can be generated in the absence of GlyA, presumably from pyruvate via serine dehydratase.

We next tried to generate a Δ*glyA* Δ*gcvPAB* double deletion in the *fhs^-^* EGD-e background, but all attempts failed, suggesting that at least one pathway for 1C-THF generation must be present for viability. Because of this observation, deletion of *glyA* and *gcvPAB* was tried in the EGD-e *fhs*^+^ background, which turned out to be possible. The resulting *fhs^+^* Δ*glyA* Δ*gcvPAB* strain exhibited wildtype growth in BHI medium, but was unable to grow in synthetic medium (Fig. 6A-B). Thus, Fhs activity is sufficient to maintain growth in complex medium but not in synthetic medium. Following this, an *fhs^-^* Δ*glyA* i*gcvPAB* mutant was generated in the EGD-e background. This strain lacks GlyA and Fhs activity and GCS activity is IPTG-dependent. This mutant could barely grow without IPTG even in BHI broth, while normal growth was observed with IPTG (Fig. 6A). Thus, at least one pathway for 1C-THF generation is essential for growth.

**Figure 6:**
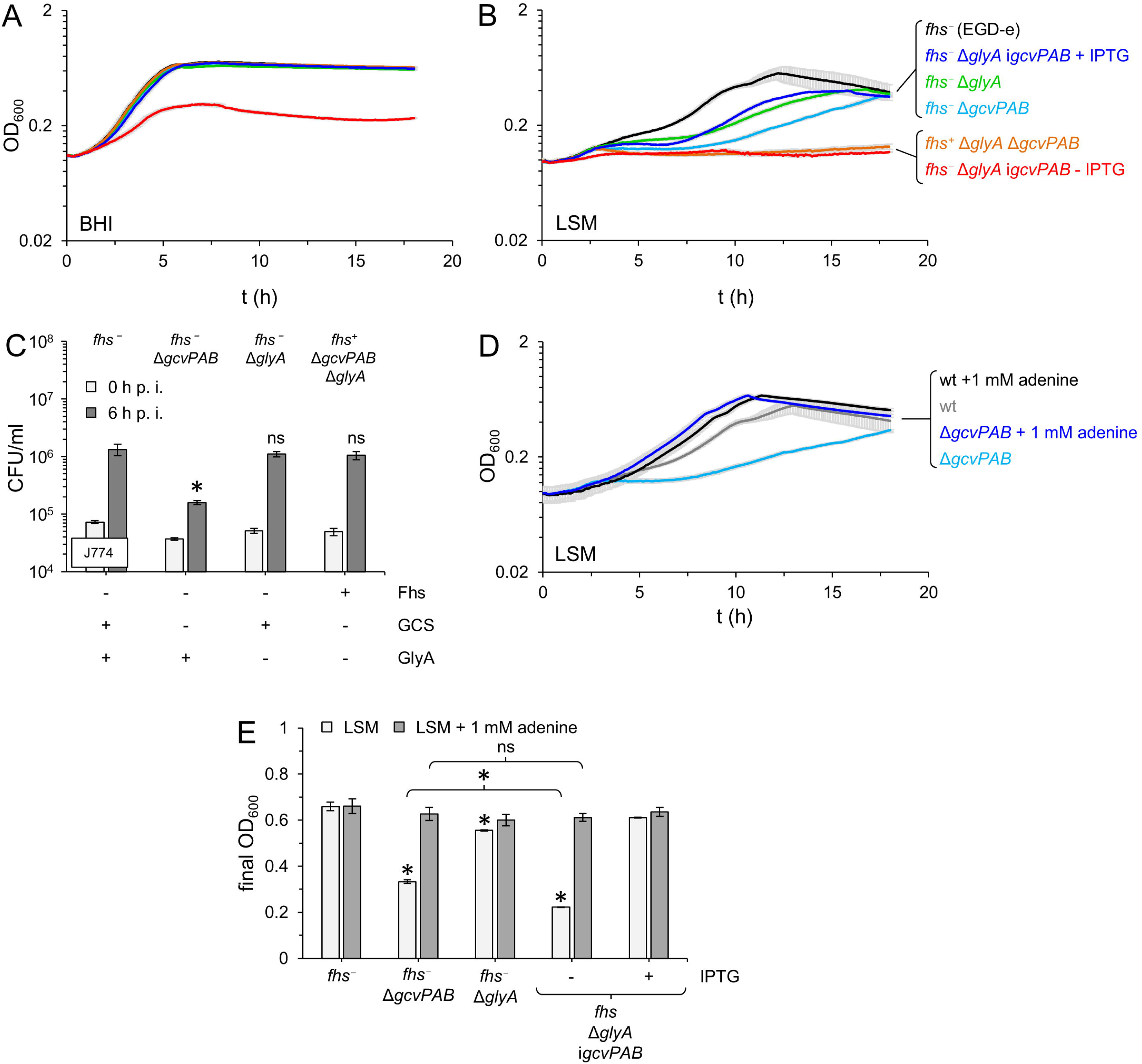
Importance of the GCS, GlyA and Fhs for viability, intracellular growth and adenine biosynthesis. (A-B) Simultaneous absence of the GCS, GlyA and Fhs activity is lethal. Growth of *L. monocyctogenes* strains EGD-e (*fhs^-^*), LMS305 (*fhs^-^* Δ*gcvPAB*), LMSF25 (*fhs^-^* Δ*glyA*), LMTE151 (*fhs^+^* Δ*gcvPAB* Δ*glyA*) and LMSF28 (*fhs^-^*Δ*glyA* i*gcvPAB*) in BHI (A) and LSM broth (B) containing or not containing 1 mM IPTG. Growth measurements were repeated three times, with each run consisting of technical replicates (n = 3). Mean values and standard deviations were calculated from the technical replicates of a representative run. (C) Individual contribution of the three 1C-THF generating pathways to intracellular growth in macrophages. Multiplication of *L. monocyctogenes* strains EGD-e (*fhs^-^*), LMS305 (*fhs^-^* Δ*gcvPAB*), LMSF25 (*fhs^-^* Δ*glyA*), LMTE151 (*fhs^+^* Δ*gcvPAB* Δ*glyA*) inside J774 mouse macrophages within six hours post infection. Infection experiments were repeated three times, with each repetition consisting of three technical replicates. Mean values and standard deviations were calculated from the technical replicates of a representative run. Statistical significance is labelled by an asterisk (*P*<0.01 *t*-test with Bonferroni-Holm correction) or “ns” (not significant). The presence or absence of the three pathways is indicated below the diagram. (D) Growth of *L. monocytogenes* strains EGD-e (wt) and LMS305 (Δ*gcvPAB*) in LSM containing standard (18 µM) and increased adenine concentrations (1 mM). Growth measurements were repeated three times, with each run consisting of technical replicates (n = 3). Mean values and standard deviations were calculated from the technical replicates of a representative run. (E) Complementation of growth defects of mutants lacking 1C-THF generating pathways by adenine supplementation. *L. monocytogenes* strains EGD-e (*fhs^-^*), LMS305 (*fhs^-^* Δ*gcvPAB*), LMSF25 (*fhs^-^* Δ*glyA*) and LMSF28 (*fhs^-^* Δ*glyA* i*gcvPAB*) were grown in LSM broth and LSM broth supplemented with 1 mM adenine. LMSF28 cultures were also grown ± 1 mM IPTG. Final optical densities were determined after 18 hours of growth. OD measurements were repeated three times, with each run consisting of technical replicates (n = 3). Mean values and standard deviations were calculated from the technical replicates of a representative run. Asterisks mark statistically significant differences (*P*<0.01, *t*-test with Bonferroni-Holm correction where necessary, ns – not significant).

Next, the individual contribution of the three 1C-THF forming pathways to intracellular growth in macrophages was measured. Of the three mutants that still possessed only one of the three biosynthetic pathways, normal intracellular proliferation was observed only in mutants that still possessed either the GCS or the Fhs/FolD pathway (Fig. 6C).

It has been shown that mutants with 1C-THF biosynthesis defects are auxotrophic for purines (Feng *et al*., 2023). We therefore tested whether addition of adenine could restore growth of Δ*gcvPAB,* mutant. As can be seen in Fig. 6D, addition of excess adenine restored normal growth of the Δ*gcvPAB* mutant, demonstrating that reduced 1C-THF biosynthesis that occurs in the absence of a functional GCS causes partial purine auxotrophy. Remarkably, excess adenine even restored growth of the *fhs^-^* Δ*glyA* i*gcvPAB* mutant in LSM broth lacking IPTG (Fig. 6E), demonstrating that the synthetic lethality of *gcvPAB*, *glyA*, and *fhs* can be attributed to their critical role in 1C-THF biosynthesis, which is required for purine formation. Quantification of 1C-THF compounds in cell extracts is technically challenging. Therefore, we opted to test the synthetic lethality of *gcvPAB* and *glyA* with *folD*, the second gene in the Fhs/FolD pathway, as a means to provide further support for this hypothesis. The *folD* gene is essential in EGD-e (Fischer *et al*., 2022), most likely explained by *fhs* inactivation. However, we were able to delete *folD* in an EGD-e background carrying a reconstituted *fhs* gene and the resulting *fhs*^+^ Δ*folD* strain was as viable as a *fhs*^+^ Δ*glyA* Δ*gcvPAB* strain (Fig. 7A). However, a *fhs*⁺ Δ*folD* Δ*glyA* i*gcvPAB* strain required IPTG for growth in BHI medium (Fig. 7A), indicating that the simultaneous deletion of *glyA* and *gcvPAB* is not tolerated in the absence of *folD*, similar to what is observed in the absence of *fhs* (see above). Notably, this growth defect was not rescued by adenine supplementation but was alleviated by thymine addition during growth in LSM medium, consistent with the metabolic model shown in Fig. 1A (Fig. 7B).

**Figure 7:**
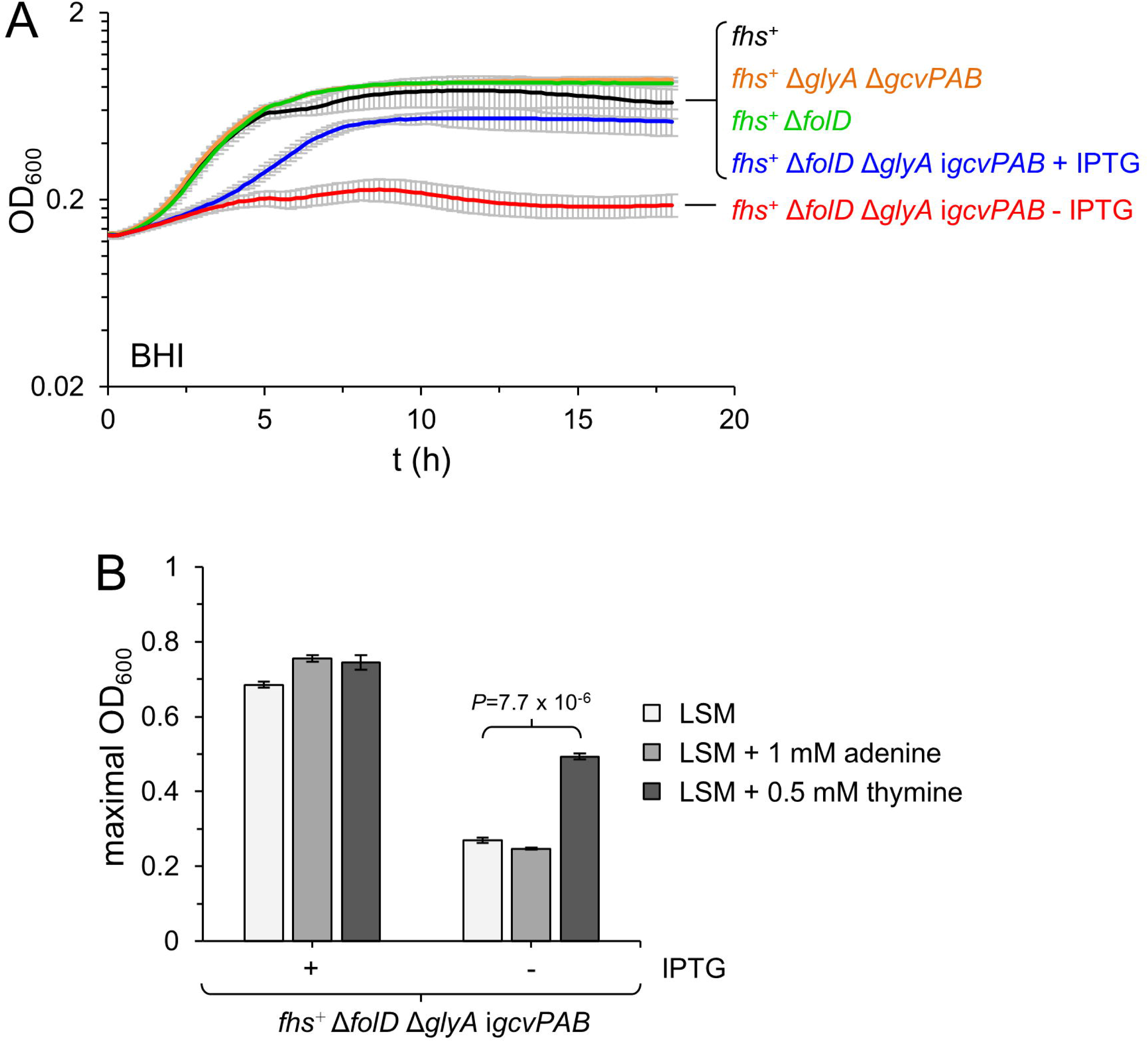
Synthetic lethality of the GCS, GlyA and FolD can be compensated by thymine addition. (A) Simultaneous absence of the GCS, GlyA and FolD activity is lethal. Growth of *L. monocytogenes* strains LMSF26 (*fhs^+^*), LMTE151 (*fhs^+^* Δ*gcvPAB* Δ*glyA*), LMMM14 (*fhs^+^* Δ*folD*) and LMMM13 (*fhs^+^* Δ*folD* Δ*glyA* i*gcvPAB*) in BHI broth ± 1 mM IPTG. For strain LMMM13, pre-depleted cells were used as inoculum. (B) Thymine but not adenine addition rescues growth of strain LMMM13 (*fhs^+^* Δ*folD* Δ*glyA* i*gcvPAB*) in the absence of IPTG. Strain LMMM13 was cultivated in LSM broth ± 1 mM IPTG and grown at 37°C for 24 h. The same experiment was carried out in LSM broth containing 1 mM adenine or 0.5 mM thymine and the maximal optical density was determined. For strain LMMM13, cells grown in the presence of IPTG were used as inoculum. Both growth measurements were repeated three times, with each run consisting of technical replicates (n = 3). Mean values and standard deviations were calculated from the technical replicates of a representative run. The *P*-value (*t*-test with Bonferroni-Holm correction) for the most relevant comparison is indicated.

## Discussion

Here we have elucidated the cause of the virulence defect of the *L. monocytogenes* Δ*gcvPAB* mutant, which shows delayed intracellular growth in the cytoplasm of macrophages and fibroblasts, the latter also explaining the spreading defect. Mice infected with this mutant lose less weight than the wildtype during a nine day period, even though statistically significant effects on bacterial replication were not observed at day three post infection. Due to the lack of the two glycine decarboxylase subunits GcvPA and GcvPB, the glycine cleavage system (GCS) is not functional in the Δ*gcvPAB* mutant. This system feeds the 1C-THF pool and a shortage of one carbon units needed for several anabolic reactions is the reason for the attenuated phenotype of the Δ*gcvPAB* strain. The observation that reactivation of the naturally inactive *fhs* gene reverses the phenotype of the Δ*gcvPAB* mutant is the most important piece of evidence for this conclusion. The *fhs* gene is split into two fragments by a premature stop codon in *L. monocytogenes* EGD-e. Formate tetrahydrofolate ligases are proteins with three domains. Their active center is located in the larger first domain (domain A), while domains B and C are probably used for oligomerisation (Radfar *et al*., 2000, Kim *et al*., 2020). The premature stop codon in EGD-e *fhs* disconnects the entire third domain, most likely inactivating the protein. Either the GCS or serine hydroxymethyltransferase activity provided by GlyA is needed for viability in the *fhs^-^* EGD-e background, whereas *gcvPAB* and *glyA* together can only be deleted in a strain with a restored full-length *fhs* gene. The reactions mediated by these three pathways all replenish the 1C-THF pool and therefore we conclude that *fhs* reactivation compensates 1C-THF depletion in the Δ*gcvPAB* mutant. This effect is particularly evident in synthetic medium, where the growth defect of the Δ*gcvPAB* mutant could also be reversed by adenine supplementation, and during intracellular growth. Both observations together point towards a limited availability of folates or folate-depending metabolites such as adenine in the host cell cytoplasm.

That glycine is toxic for the Δ*gcvPAB* mutant is another argument for the 1C-THF depletion hypothesis: 1C-THF cannot be generated from glycine in the Δ*gcvPAB* mutant and 1C-THF formation by GlyA is prevented in the presence of excess glycine at the same time. This is because the GlyA-mediated reaction is reversible (Schirch & Szebenyi, 2005) and therefore shifted towards the 1C-THF consuming formation of serine (Fig. 1A). The reactivation of Fhs partially neutralizes the toxic effect of glycine, as this counteracts the 1C-THF depletion resulting from increased serine biosynthesis. Limitations in 1C-THF availability also explains the contradiction between the reported essentiality of the *folD* gene in EGD-e (Fischer *et al*., 2022) and the successful *folD* deletion in strain 10403S (Feng *et al*., 2023): N10-formyl-THF, a 1C-THF species needed for purine and N-formylmethionine biosynthesis can only be generated by Fhs or FolD and therefore, a *fhs folD* double mutant has pronounced growth defects in laboratory media and during infection (Feng *et al*., 2023). That is why transposon insertion mutants in the *folD* gene were likely counter-selected in our recent Tn-Seq study that was performed in the *fhs^-^*EGD-e background (Fischer *et al*., 2022).

Two enzymatic steps in THF biosynthesis ahead of Fhs are inhibited by cotrimoxazole (Fig. 1A), an antibiotic that is recommended for the treatment of listeriosis (Karsaliakos & Mylonakis, 2023). Our experiments revealed that collective inactivation of the Fhs/FolD pathway, the GCS and GlyA strongly impaired growth, suggesting the absence of other 1C-THF generating pathways in *L. monocytogenes*. Therefore, it would be interesting to test whether inhibitors of any of these three pathways such as the pyrazolopyran compounds inhibiting serine hydroxymethyltransferase (Makino *et al*., 2022) would act synergistically with cotrimoxazole.

A certain hierarchy of 1C-THF-generating pathways can also be derived from our data. While each of the three pathways was sufficient to maintain growth in BHI medium (Fig. 6A), the GCS and GlyA had a greater impact on growth in synthetic medium than Fhs/FolD (Fig. 6B). In contrast, the GCS or Fhs/FolD were each sufficient to maintain growth in macrophages. Thus, the GCS is important for 1C-THF formation under all tested conditions, whereas GlyA and the Fhs/FolD are required under specific conditions only and are therefore of secondary importance.

Several genes frequently inactivated by premature stop codons are known in reference strains and isolates of *L. monocytogenes*, including the internalin gene *inlA* or the *gadR* acid resistance regulator gene (Nightingale et al., 2005, Wu et al., 2023). As far as we know, the *fhs* gene is the first example of such a cleaved and inactive pseudogene that can be reactivated by suppressor mutations if the selection conditions favor its restoration. This proves that the inactivation of genes by premature stop codons is not an evolutionary dead end, but can be a reversible regulatory event of transient nature. It would be important to find out whether similar effects can also be observed on *inlA* genes inactivated by premature stop codons, because this would have important implications on the risk assessment of strains with such mutations.

## Materials and Methods

### Bacterial strains and growth conditions

All *L. monocytogenes* strains are listed in Table 1. Strains were generally cultivated in BHI broth or on BHI agar plates. LSM broth and LSM agar plates were used for cultivation under chemically defined conditions (Whiteley *et al*., 2017). Antibiotics and supplements were added at the following concentrations: erythromycin (5 µg/ml), kanamycin (50 µg/ml), X-Gal (100 µg/ml) and IPTG (1 mM). *Escherichia coli* TOP10 was used as standard cloning host (Sambrook *et al*., 1989).

**Table 1:**
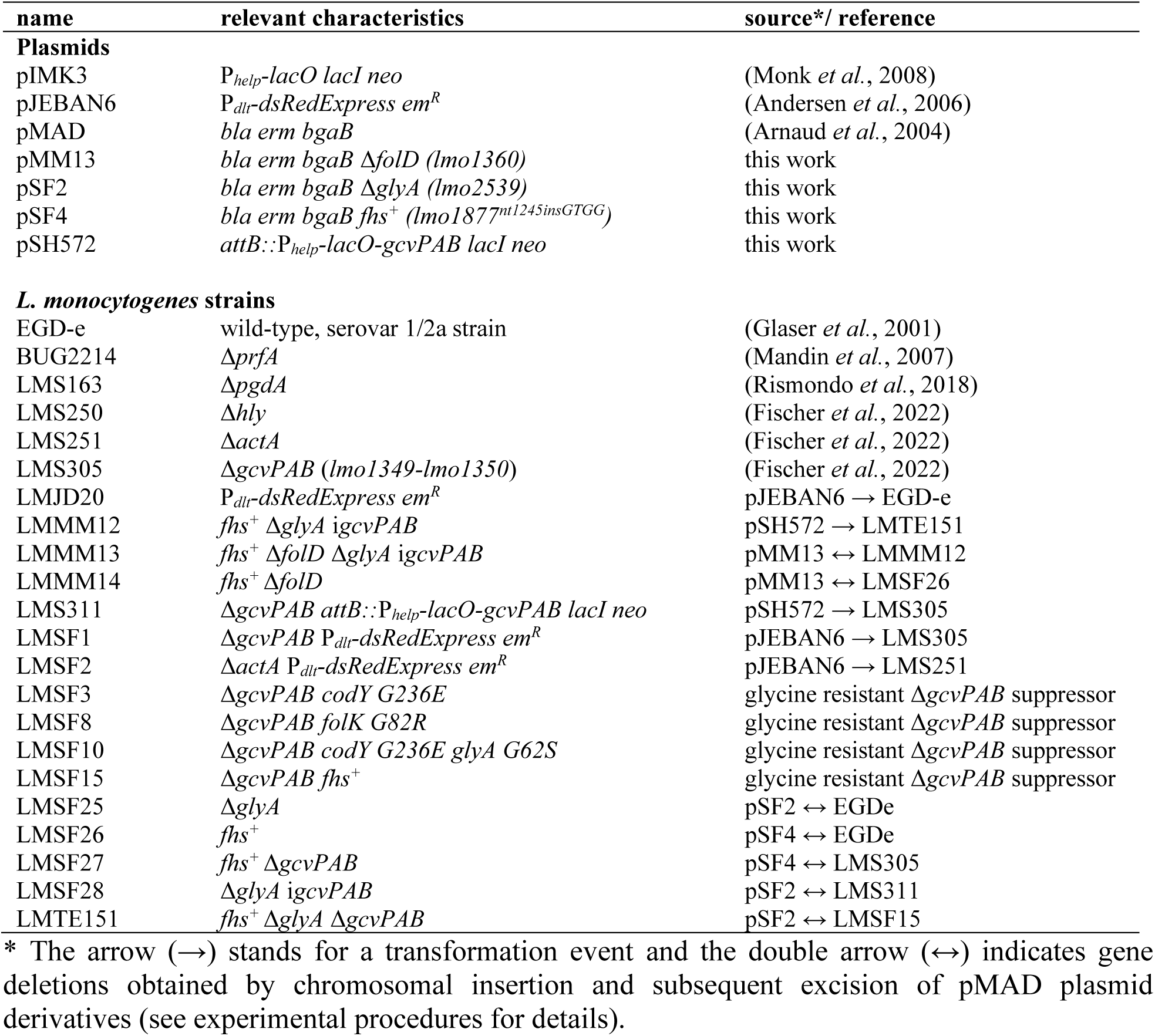
Plasmids and *L. monocytogenes* strains used in this study.

### General methods, manipulation of DNA and oligonucleotide primers

Standard methods were used for transformation of *E. coli* and isolation of plasmid DNA (Sambrook *et al*., 1989). Transformation of *L. monocytogenes* was carried out as described by others (Monk *et al*., 2008). Restriction and ligation of DNA was performed according to the manufactureŕs instructions. The oligonucleotides used in this study are listed in Table 2.

**Table 2:**
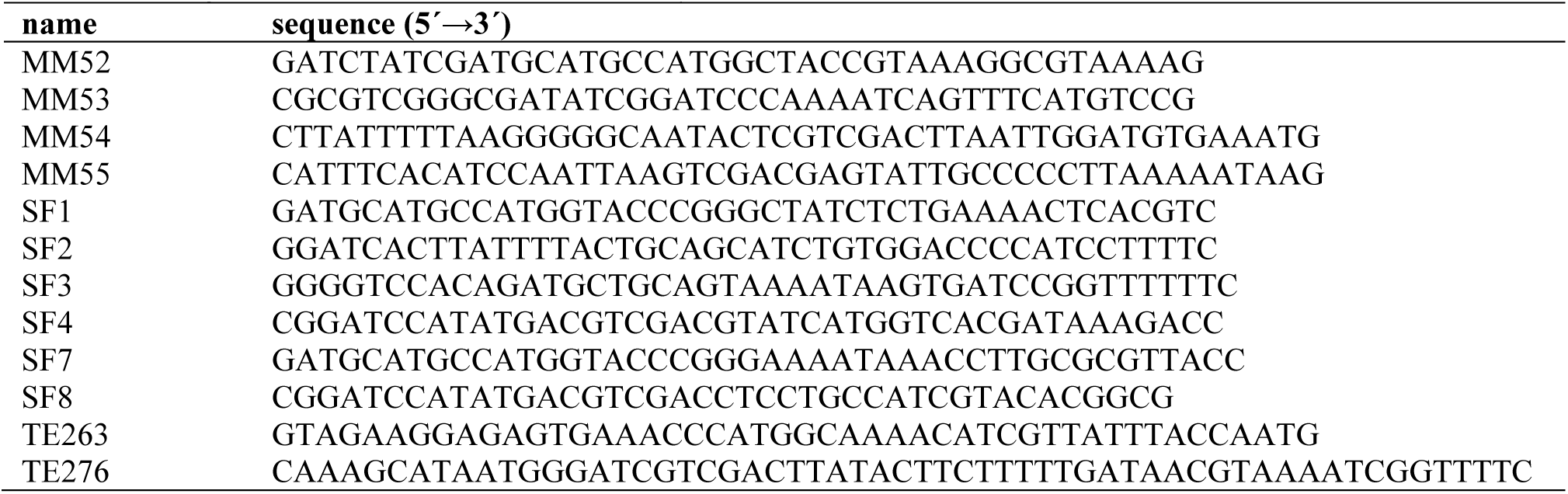
Oligonucleotides used in this study.

### Construction of plasmids and *L. monocytogenes* strains

All plasmids are listed in Table 1. Plasmid pSF2 was generated for deletion of *glyA*. To this end, fragments up-and downstream of *glyA* were amplified by PCR using SF1/SF2 and SF3/SF4 as the primers, respectively. The resulting fragments were fused together by splicing by overlapping extension (SOE) PCR and then inserted into pMAD using NcoI/SalI as the restriction enzymes. Plasmid pMM13 was constructed for *folD* deletion. Fragments up-and downstream of *folD* were amplified from chromosomal DNA using the oligonucleotide pairs MM52/MM55 and MM54/MM53, respectively. Both fragments were joined by SOE-PCR and inserted into pMAD by restriction-free cloning.

Plasmid pSF4 was constructed to transfer the *fhs*^+^ mutation of strain LMSF15 to other strain backgrounds. For this, the *fhs*^+^ region of strain LMSF15 was amplified using SF7/SF8 as the primers and the obtained fragment was inserted into pMAD using NcoI/SalI. The plasmid insertion/excision protocol of Arnaud et al. (Arnaud *et al*., 2004) was used for gene deletions and allelic exchange. Successful deletions and allelic exchanges were confirmed by PCR and genome sequencing (see below).

Plasmid pSH572 was generated for complementation of the Δ*gvcPAB* mutant. To this end, the *gcvPAB* operon was amplified from EGD-e chromosomal DNA in a PCR using the oligonucleotides TE263/TE276. The resulting fragment was cut using NcoI/SalI and ligated to pIMK3 digested with the same enzymes. Plasmid pSH572 was transformed into strain LMS305 and kanamycin resistant clones were selected. Plasmid insertion at the *attB* tRNA^Arg^ site was confirmed by PCR.

pJEBAN6 for expression of DsRed-Express was electroporated in various recipient strains and erythromycin resistant clones were selected. Plasmid acquisition and concomitant DsRed-Express production was confirmed by fluorescence microscopy.

### Genome sequencing

Chromosomal DNA of bacterial strains was isolated using the GenElute Bacterial Genomic DNA Kit (Sigma-Aldrich). Libraries were generated from 1 ng DNA using the Nextera XT DNA Library Prep Kit (Illumina). Sequencing was carried out on a NextSeq sequencer in paired-end sequencing mode with 2 x 150 bp read length. Reads were mapped to the *L. monocytogenes* EGD-e genome (NC_003210.1) (Glaser *et al*., 2001) as the reference in Geneious (Biomatters Ltd.) and the alignment was analyzed using the Geneious SNP finder tool. Genome sequences of the Δ*gcvPAB* mutant and glycine resistant Δ*gcvPAB* suppressors were deposited at the European Nucleotide Archive (ENA, https://www.ebi.ac.uk/) under project accession number PRJEB94141.

### Hemolysis assays

The CAMP test was used as a qualitative assay to record hemolysis (Fernandez-Garayzabal *et al*., 1996). For this, *Staphylococcus aureus* SG511 was streaked on Mueller-Hinton agar plates containing 5% sheep blood right-angled to the *L. monocytogenes* strains to be tested. Hemolysis became apparent after overnight incubation at 37°C.

For semi-quantitative determination of hemolysis activity, bacterial cultures were cultivated for 5 h at 37°C. Culture supernatants were prepared from these cultures by centrifugation of a 1 ml culture volume and a two-fold dilution series of the culture supernatant was prepared in phosphate-buffered saline (PBS) containing 6 mM cysteine. Triplicates of each dilution (100 µl) were pipetted into a 96-well microtiter plate and 1 µl of a 1% (v/v) human erythrocyte concentrate was added to each well. The microtiter plate was incubated for 30 min at 37°C in a static incubator. Hemolytic activity is expressed as the dilution factor of the last dilution showing complete hemolysis.

### Assays to determine hydrogen peroxide and lysozyme sensitivity

To assess the sensitivity of *L. monocytogenes* to H2O2, strains were inoculated in BHI broth containing increasing concentrations of H2O2 (0.2-100 mM) in a microtiter plate at 37°C. The plate was incubated overnight at 37 °C in a static incubator. The minimum inhibitory concentration of hydrogen peroxide was determined the next morning by visual inspection.

For analysis of lysozyme sensitivity, *L. monocytogenes* strains were grown in BHI broth at 37°C until mid-logarithmic growth phase (OD600∼0.8). Cells were collected by centrifugation and resuspended in 50 mM Tris/HCl pH8.0 to an optical density of OD600=0.6. Lysozyme was added to a final concentration of 2.5 µg/ml and the cells were shaken at 37°C. Lysis was followed by measuring the decrease in optical density (λ=600 nm) every 15 min in a spectrophotometer.

### Cell culture infection experiments

Infection of J774A.1 mouse macrophages (ATCC® TIB-67^TM^) and HepG2 human hepatocytes (ATCC® HB-8065^TM^) with *L. monocytogenes* strains was performed as described earlier (Halbedel *et al*., 2019). Briefly, 10^5^ cells were seeded into the wells of a 24 multi well plate and cultivated in DMEM + 10% fetal calf serum (FCS) overnight before they were infected with 2 x 10^5^ bacteria. The bacteria were allowed to invade the cells during an incubation step at 37°C for 1 h. Extracellular bacteria were first washed off with PBS and the remaining extracellular bacteria were killed by gentamicin addition. Eukaryotic cells were lysed six hours post infection using ice-cold PBS containing 0.1% Triton X-100, serial dilutions were plated on BHI agar plates and incubated overnight at 37°C for quantification. Infection of 3T3-L1 mouse embryo fibroblasts (ATCC® CL-173T^M^) was performed in the same way. Analysis of cell-to-cell spread using 3T3-L1 mouse embryo fibroblasts by plaque formation was carried out as described earlier (Halbedel *et al*., 2014). Here, 5 x 10^5^ fibroblast cells were seeded into the wells of a six well plate and cultivated in DMEM + 10% newborn calf serum (NCS). After three days of incubation, cells were infected with an inoculum of 2, 4 or 10 μl each containing 1 x 10^6^ bacteria. Plaques were visualised three days post infection using neutral red staining. Plaque areas were quantified using ImageJ.

### Microscopy of infected 3T3 cells

5 x 10^4^ 3T3 mouse fibroblasts were seeded into the wells of a 24-well tissue culture plate containing coverslips and DMEM + 10% NCS as the culture medium and incubated in a 5% CO2 atmosphere at 37°C. Cells were infected as outlined above. 6 h after infection, cells were washed with PBS and then fixed with ice-cold methanol. Methanol was replaced by 500 μl PBS before the coverslips were removed from the wells and allowed to dry completely. Finally, a drop of ProLong Gold antifade reagent with DAPI (Invitrogen) was dropped onto a microscope slide and the coverslip was placed on top. Samples were dried overnight and examined using a Nikon Eclipse Ti fluorescent microscope.

### *In vivo* infection in mice

80 male C57BL/6J mice (Janvier) were maintained in the animal facility at the university hospital of the Otto-von-Guericke University of Magdeburg under conditions that ensured precise temperature and humidity regulation, with a 12-hour day/night cycle. Mice at an age ranging from 10 to 18 weeks were infected with 1-20 x 10^4^ CFU in PBS via the tail vein. Following infection, the mice were monitored daily for body weight loss and other disease symptoms for the entire duration of the experiment. After euthanasia of mice by carbon dioxide (CO2) inhalation, the heart was punctured, the heart blood was taken and the heart was perfused with 10 mL PBS. All animal experiments were conducted according to the institutional guidelines, and the study was approved by local government agencies (Landesverwaltungsamt Sachsen-Anhalt; AZ 42502-2-1603 UniMD).

### Determination of CFU in spleen, brain and liver

To determine the CFUs in the spleen, brain and liver, the organs were homogenised in 0.2% IGEPAL CA-630 (Sigma-Aldrich) lysis buffer. The livers were suspended in 2 ml IGEPAL buffer, while the brains and spleens were suspended in 1 ml IGEPAL buffer. The organs were homogenised at full speed (30000 rpm) using a POLYTRON® PT 3100 homogeniser (KINEMATICA AG). To prevent artefacts between samples, the homogeniser was washed with EtOH for 10 seconds three times and with PBS twice between every sample. Serial dilutions were plated onto BHI agar plates and incubated at 37 °C for 24 hours to quantify the colonies.

## Acknowledgements

We thank Birgitt Hahn for excellent technical assistance. We also thank Nouria Jantz-Naeem, Anna Krone, Bianca Thoma, Hildburg Volkmann, Anne Hoffmann and Anja Sammt from the Kahlfuß lab for helping processing the samples following the *in vivo* infection model. This work was funded by the DFG (grants HA6830/2-1 and HA6830/5-1 to S. H.).

## References

Andersen, J.B., Roldgaard, B.B., Lindner, A.B., Christensen, B.B., and Licht, T.R. (2006) Construction of a multiple fluorescence labelling system for use in co-invasion studies of *Listeria monocytogenes*. BMC Microbiol 6: 86, 10.1186/1471-2180-6-86.

Arnaud, M., Chastanet, A., and Debarbouille, M. (2004) New vector for efficient allelic replacement in naturally nontransformable, low-GC-content, gram-positive bacteria. Appl Environ Microbiol 70: 6887–6891, 10.1128/AEM.70.11.6887-6891.2004.

Bagatella, S., Tavares-Gomes, L., and Oevermann, A. (2022) *Listeria monocytogenes* at the interface between ruminants and humans: A comparative pathology and pathogenesis review. Vet Pathol 59: 186–210, 10.1177/03009858211052659.

Borezee, E., Pellegrini, E., and Berche, P. (2000) OppA of *Listeria monocytogenes*, an oligopeptide-binding protein required for bacterial growth at low temperature and involved in intracellular survival. Infect Immun 68: 7069–7077, 10.1128/iai.68.12.7069-7077.2000.

Charlier, C., Disson, O., and Lecuit, M. (2020) Maternal-neonatal listeriosis. Virulence 11: 391–397, 10.1080/21505594.2020.1759287.

Chico-Calero, I., Suarez, M., Gonzalez-Zorn, B., Scortti, M., Slaghuis, J., Goebel, W., Vazquez-Boland, J.A., and European Listeria Genome, C. (2002) Hpt, a bacterial homolog of the microsomal glucose-6-phosphate translocase, mediates rapid intracellular proliferation in *Listeria*. Proc Natl Acad Sci U S A 99: 431–436, 10.1073/pnas.012363899.

Disson, O., and Lecuit, M. (2012) Targeting of the central nervous system by *Listeria monocytogenes*. Virulence 3: 213–221, 10.4161/viru.19586.

Eisenhut, M., Bauwe, H., and Hagemann, M. (2007) Glycine accumulation is toxic for the cyanobacterium *Synechocystis* sp. strain PCC 6803, but can be compensated by supplementation with magnesium ions. FEMS Microbiol Lett 277: 232–237, 10.1111/j.1574-6968.2007.00960.x.

Faith, N.G., Kim, J.W., Azizoglu, R., Kathariou, S., and Czuprynski, C. (2012) Purine biosynthesis mutants (*purA* and *purB*) of serotype 4b *Listeria monocytogenes* are severely attenuated for systemic infection in intragastrically inoculated A/J Mice. Foodborne Pathog Dis 9: 480–486, 10.1089/fpd.2011.1013.

Feng, Y., Chang, S.K., and Portnoy, D.A. (2023) The major role of *Listeria monocytogenes* folic acid metabolism during infection is the generation of N-formylmethionine. mBio: e0107423, 10.1128/mbio.01074-23.

Fernandez-Garayzabal, J.F., Suarez, G., Blanco, M.M., Gibello, A., and Dominguez, L. (1996) Taxonomic note: a proposal for reviewing the interpretation of the CAMP reaction between *Listeria monocytogenes* and *Rhodococcus equi*. Int J Syst Bacteriol 46: 832–834, 10.1099/00207713-46-3-832.

Fischer, M.A., Engelgeh, T., Rothe, P., Fuchs, S., Thürmer, A., and Halbedel, S. (2022) *Listeria monocytogenes* genes supporting growth under standard laboratory cultivation conditions and during macrophage infection. Genome Res 32: 1711–1726, 10.1101/gr.276747.122.

Freeman, M.J., Eral, N.J., and Sauer, J.D. (2025) *Listeria monocytogenes* requires phosphotransferase systems to facilitate intracellular growth and virulence. PLoS Pathog 21: e1012492, 10.1371/journal.ppat.1012492.

Fujiwara, K., Okamura-Ikeda, K., and Motokawa, Y. (1984) Mechanism of the glycine cleavage reaction. Further characterization of the intermediate attached to H-protein and of the reaction catalyzed by T-protein. J Biol Chem 259: 10664–10668,

Glaser, P., Frangeul, L., Buchrieser, C., Rusniok, C., Amend, A., Baquero, F., Berche, P., Bloecker, H., Brandt, P., Chakraborty, T., Charbit, A., Chetouani, F., Couve, E., de Daruvar, A., Dehoux, P., Domann, E., Dominguez-Bernal, G., Duchaud, E., Durant, L., Dussurget, O., Entian, K.D., Fsihi, H., Garcia-del Portillo, F., Garrido, P., Gautier, L., Goebel, W., Gomez-Lopez, N., Hain, T., Hauf, J., Jackson, D., Jones, L.M., Kaerst, U., Kreft, J., Kuhn, M., Kunst, F., Kurapkat, G., Madueno, E., Maitournam, A., Vicente, J.M., Ng, E., Nedjari, H., Nordsiek, G., Novella, S., de Pablos, B., Perez-Diaz, J.C., Purcell, R., Remmel, B., Rose, M., Schlueter, T., Simoes, N., Tierrez, A., Vazquez-Boland, J.A., Voss, H., Wehland, J., and Cossart, P. (2001) Comparative genomics of *Listeria* species. Science 294: 849–852, 10.1126/science.1063447294/5543/849 [pii].

Green, J.M., and Matthews, R.G. (2007) Folate Biosynthesis, Reduction, and Polyglutamylation and the Interconversion of Folate Derivatives. EcoSal Plus 2, 10.1128/ecosalplus.3.6.3.6.

Halbedel, S., Prager, R., Banerji, S., Kleta, S., Trost, E., Nishanth, G., Alles, G., Holzel, C., Schlesiger, F., Pietzka, A., Schlüter, D., and Flieger, A. (2019) A *Listeria monocytogenes* ST2 clone lacking chitinase ChiB from an outbreak of non-invasive gastroenteritis. Emerg Microbes Infect 8: 17–28, 10.1080/22221751.2018.1558960.

Halbedel, S., Reiss, S., Hahn, B., Albrecht, D., Mannala, G.K., Chakraborty, T., Hain, T., Engelmann, S., and Flieger, A. (2014) A systematic proteomic analysis of *Listeria monocytogenes* house-keeping protein secretion systems. Molecular & cellular proteomics: MCP 13: 3063–3081, 10.1074/mcp.M114.041327.

Karsaliakos, P.M., and Mylonakis, E. 2023. Listeriosis - symptoms, diagnosis and treatment. https://bestpractice.bmj.com/topics/en-us/914

Kikuchi, G., Motokawa, Y., Yoshida, T., and Hiraga, K. (2008) Glycine cleavage system: reaction mechanism, physiological significance, and hyperglycinemia. Proc Jpn Acad Ser B Phys Biol Sci 84: 246–263, 10.2183/pjab.84.246.

Kim, S., Lee, S.H., Seo, H., and Kim, K.J. (2020) Biochemical properties and crystal structure of formate-tetrahydrofolate ligase from *Methylobacterium extorquens* CM4. Biochemical and biophysical research communications 528: 426–431, 10.1016/j.bbrc.2020.05.198.

Kocks, C., Gouin, E., Tabouret, M., Berche, P., Ohayon, H., and Cossart, P. (1992) *L. monocytogenes*-induced actin assembly requires the *actA* gene product, a surface protein. Cell 68: 521–531, 10.1016/0092-8674(92)90188-i.

Koopmans, M.M., Brouwer, M.C., Vazquez-Boland, J.A., and van de Beek, D. (2023) Human Listeriosis. Clinical microbiology reviews 36: e0006019, 10.1128/cmr.00060-19.

Makino, Y., Oe, C., Iwama, K., Suzuki, S., Nishiyama, A., Hasegawa, K., Okuda, H., Hirata, K., Ueno, M., Kawaji, K., Sasano, M., Usui, E., Hosaka, T., Yabuki, Y., Shirouzu, M., Katsumi, M., Murayama, K., Hayashi, H., and Kodama, E.N. (2022) Serine hydroxymethyltransferase as a potential target of antibacterial agents acting synergistically with one-carbon metabolism-related inhibitors. Commun Biol 5: 619, 10.1038/s42003-022-03555-x.

Mandin, P., Repoila, F., Vergassola, M., Geissmann, T., and Cossart, P. (2007) Identification of new noncoding RNAs in *Listeria monocytogenes* and prediction of mRNA targets. Nucleic Acids Res 35: 962–974, 10.1093/nar/gkl1096.

Monk, I.R., Gahan, C.G., and Hill, C. (2008) Tools for functional postgenomic analysis of *Listeria monocytogenes*. Appl Environ Microbiol 74: 3921–3934, AEM.00314-08 [pii] 10.1128/AEM.00314-08.

Nightingale, K.K., Windham, K., Martin, K.E., Yeung, M., and Wiedmann, M. (2005) Select *Listeria monocytogenes* subtypes commonly found in foods carry distinct nonsense mutations in inlA, leading to expression of truncated and secreted internalin A, and are associated with a reduced invasion phenotype for human intestinal epithelial cells. Appl Environ Microbiol 71: 8764–8772, 10.1128/AEM.71.12.8764-8772.2005.

Pizarro-Cerda, J., and Cossart, P. (2018) *Listeria monocytogenes*: cell biology of invasion and intracellular growth. Microbiol Spectr 6, 10.1128/microbiolspec.GPP3-0013-2018.

Quereda, J.J., Moron-Garcia, A., Palacios-Gorba, C., Dessaux, C., Garcia-Del Portillo, F., Pucciarelli, M.G., and Ortega, A.D. (2021) Pathogenicity and virulence of *Listeria monocytogenes*: A trip from environmental to medical microbiology. Virulence 12: 2509–2545, 10.1080/21505594.2021.1975526.

Radfar, R., Shin, R., Sheldrick, G.M., Minor, W., Lovell, C.R., Odom, J.D., Dunlap, R.B., and Lebioda, L. (2000) The crystal structure of N(10)-formyltetrahydrofolate synthetase from *Moorella thermoacetica*. Biochemistry 39: 3920–3926, 10.1021/bi992790z.

Rismondo, J., Wamp, S., Aldridge, C., Vollmer, W., and Halbedel, S. (2018) Stimulation of PgdA-dependent peptidoglycan N-deacetylation by GpsB-PBP A1 in *Listeria monocytogenes*. Mol Microbiol 107: 472–487, 10.1111/mmi.13893.

Sambrook, J., Fritsch, E.F., and Maniatis, T., (1989) Molecular cloning: a laboratory manual, p. 3 v. Cold Spring Harbor Laboratory Press, Cold Spring Harbor, N.Y.

Schirch, V., Hopkins, S., Villar, E., and Angelaccio, S. (1985) Serine hydroxymethyltransferase from *Escherichia coli*: purification and properties. J Bacteriol 163: 1–7, 10.1128/jb.163.1.1-7.1985.

Schirch, V., and Szebenyi, D.M. (2005) Serine hydroxymethyltransferase revisited. Curr Opin Chem Biol 9: 482–487, 10.1016/j.cbpa.2005.08.017.

Smith, H.B., Li, T.L., Liao, M.K., Chen, G.Y., Guo, Z., and Sauer, J.D. (2021) *Listeria monocytogene*s MenI Encodes a DHNA-CoA Thioesterase Necessary for Menaquinone Biosynthesis, Cytosolic Survival, and Virulence. Infect Immun 89, 10.1128/IAI.00792-20.

Stamm, C.E., McFarland, A.P., Locke, M.N., Tabakh, H., Tang, Q., Thomason, M.K., and Woodward, J.J. (2024) RECON gene disruption enhances host resistance to enable genome-wide evaluation of intracellular pathogen fitness during infection. mBio 15: e0133224, 10.1128/mbio.01332-24.

Stritzker, J., Janda, J., Schoen, C., Taupp, M., Pilgrim, S., Gentschev, I., Schreier, P., Geginat, G., and Goebel, W. (2004) Growth, virulence, and immunogenicity of *Listeria monocytogenes* aro mutants. Infect Immun 72: 5622–5629, 10.1128/IAI.72.10.5622-5629.2004.

Whiteley, A.T., Garelis, N.E., Peterson, B.N., Choi, P.H., Tong, L., Woodward, J.J., and Portnoy, D.A. (2017) c-di-AMP modulates *Listeria monocytogenes* central metabolism to regulate growth, antibiotic resistance and osmoregulation. Mol Microbiol 104: 212–233, 10.1111/mmi.13622.

Wu, J., McAuliffe, O., and O’Byrne, C.P. (2023) A novel RofA-family transcriptional regulator, GadR, controls the development of acid resistance in *Listeria monocytogenes*. mBio 14: e0171623, 10.1128/mbio.01716-23.

Xayarath, B., Marquis, H., Port, G.C., and Freitag, N.E. (2009) Listeria monocytogenes CtaP is a multifunctional cysteine transport-associated protein required for bacterial pathogenesis. Mol Microbiol 74: 956–973, 10.1111/j.1365-2958.2009.06910.x.

Yishai, O., Bouzon, M., Doring, V., and Bar-Even, A. (2018) In Vivo Assimilation of One-Carbon via a Synthetic Reductive Glycine Pathway in *Escherichia coli*. ACS Synth Biol 7: 2023–2028, 10.1021/acssynbio.8b00131.

Zhang, Y., Anaya-Sanchez, A., and Portnoy, D.A. (2022) para-Aminobenzoic acid biosynthesis is required for *Listeria monocytogenes* growth and pathogenesis. Infect Immun 90: e0020722, 10.1128/iai.00207-22.

