## Supplementary Figures S1-S3 for "Three metabolic pathways replenishing the one-carbon pool collectively support growth and virulence of *Listeria monocytogenes*"

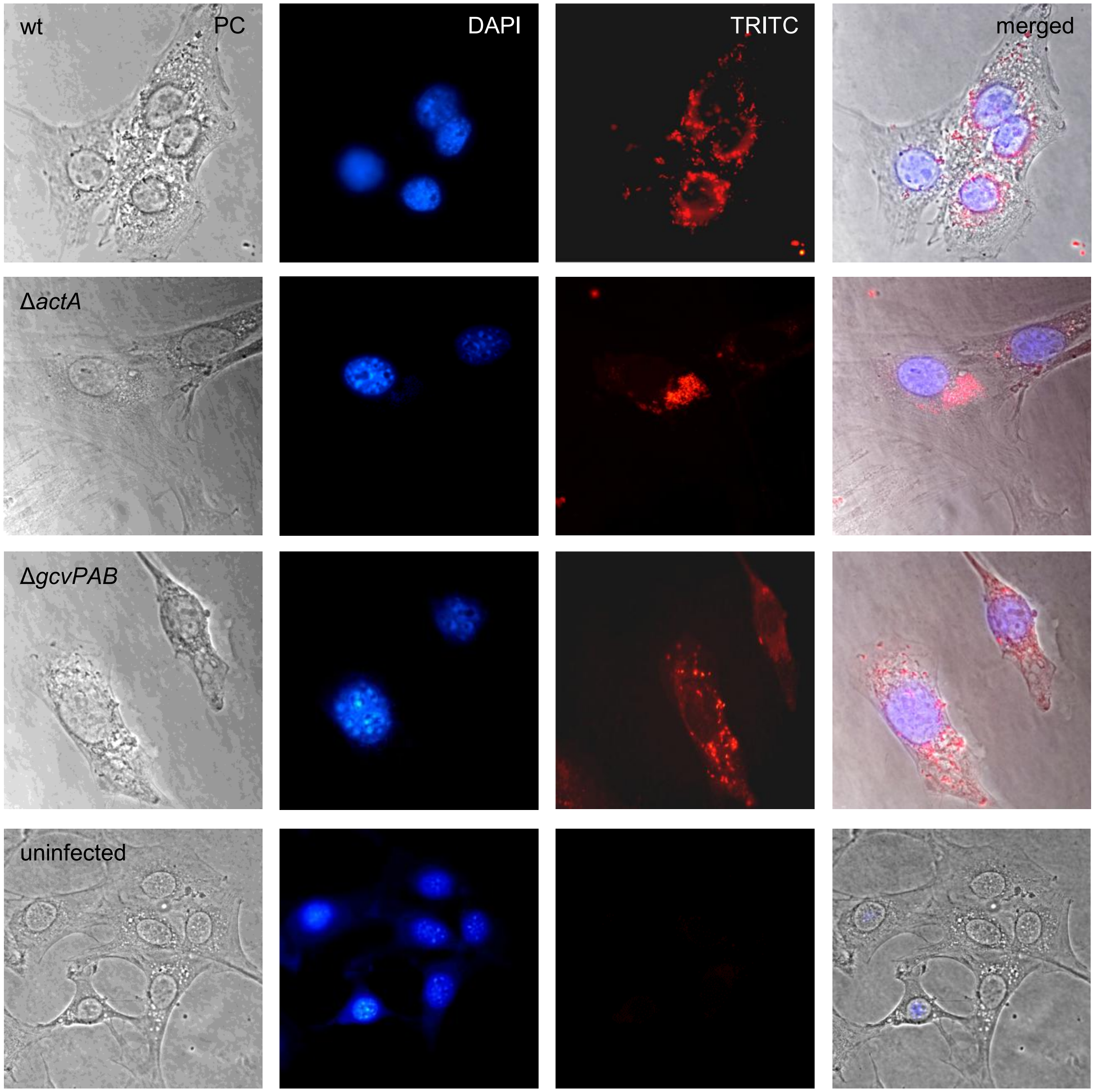


**Figure S1:** Dissemination of the Δ*gcvPAB* mutant in 3T3 mouse fibroblasts.
Micrographs showing intracellular dissemination of dsRed-Express producing *L. monocytogenes* strains LMJD20 (wt), LMFS1 (Δ*gcvPAB*) and LMSF2 (Δ*actA*) in 3T3 mouse fibroblasts six hours post infection.

Freier *et al.* Fig. S1

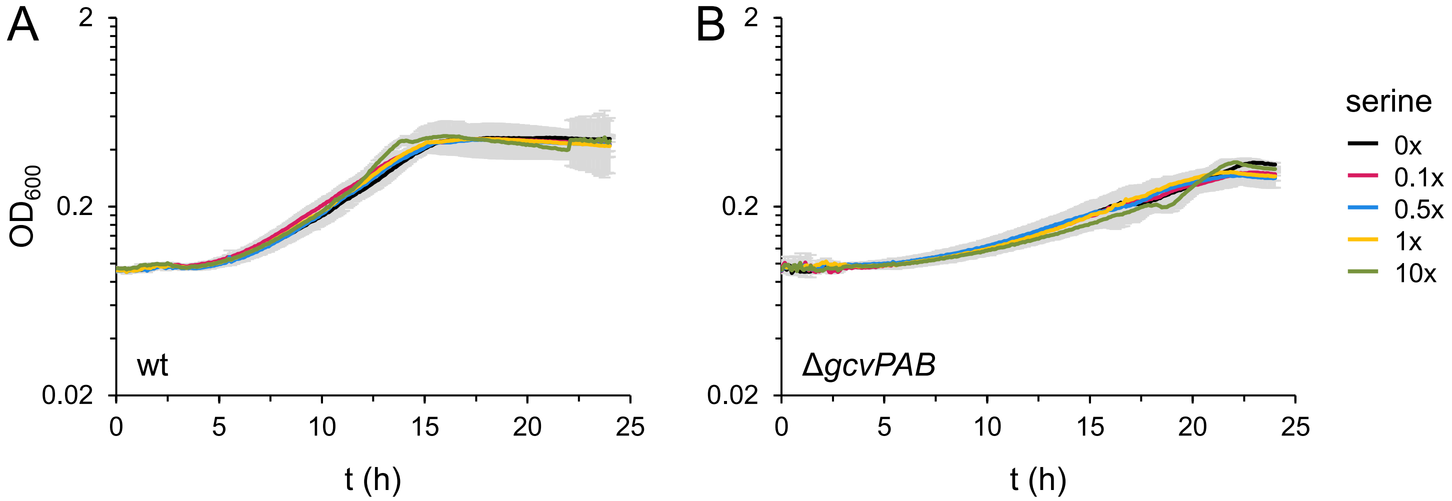


**Figure S2**: Growth of *L. monocytogenes* strains EGD-e (wt, A) and LMS305 (Δ*gcvPAB*, B) in LSM medium containing different serine concentrations. Serine concentrations are expressed relative to standard LSM concentrations (1x = 1 mM). Growth measurements were repeated three times, with each run consisting of technical replicates (n = 3). Mean values and standard deviations were calculated from the technical replicates of a representative run.

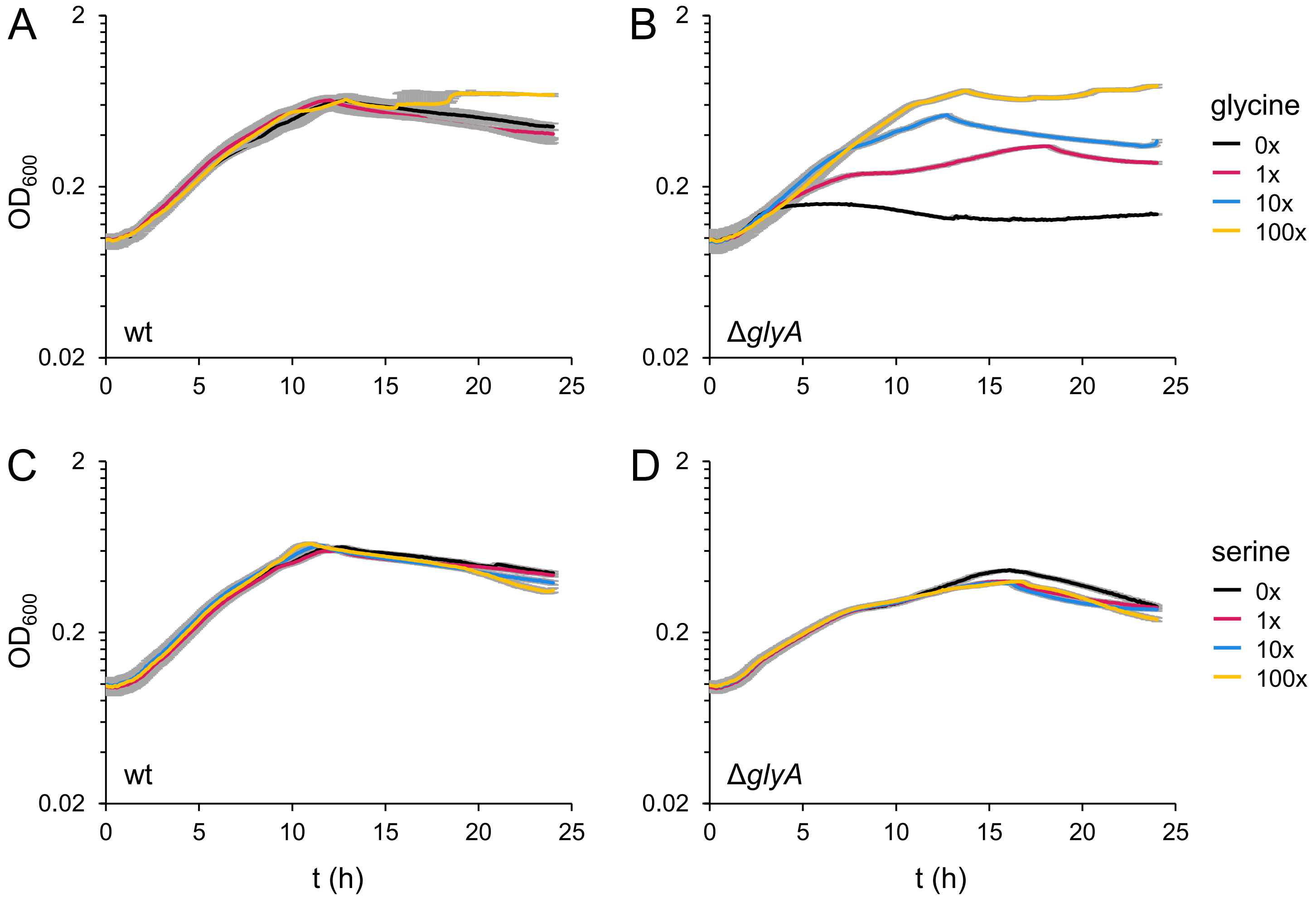


**Figure S3**: Growth of a *L. monocytogenes* Δ*glyA* mutant in LSM medium with different glycine and serine concentrations.
(A-B) Growth of *L. monocytogenes* strains EGD-e (wt, A) and LMSF25 (Δ*glyA*, B) in LSM medium containing different glycine concentrations.
(C-D) Growth of *L. monocytogenes* strains EGD-e (wt, C) and LMSF25 (Δ*glyA*, D) in LSM medium containing different serine concentrations. Glycine and serine concentrations are expressed relative to standard LSM concentrations (1x = 1.3 mM for glycine, 1x = 1 mM for serine).
Growth measurements were repeated three times, with each run consisting of technical replicates (n = 3). Mean values and standard deviations were calculated from the technical replicates of a representative run.
